# Activated gliosis, accumulation of amyloid β and hyperphosphorylation of tau in aging canines with and without cognitive decline

**DOI:** 10.1101/2022.12.21.521473

**Authors:** Amelia D. Hines, Stephanie McGrath, Amanda S. Latham, Breonna Kusick, Lisa Mulligan, McKenzie L. Richards, Julie A. Moreno

## Abstract

Canine cognitive dysfunction (CCD) syndrome is a well-recognized naturally occurring disease in aged dogs, with a remarkably similar disease course, both in its clinical presentation and neuropathological changes, as humans with Alzheimer’s disease (AD). Similar to human AD patients this naturally occurring disease is found in the aging canine population however, there is little understanding of how the canine brain ages pathologically. It is well known that in neurodegenerative diseases, there is an increase in inflamed glial cells as well as an accumulation of hyperphosphorylation of tau (P-tau) and amyloid beta (Aβ_1-42_). These pathologies increase neurotoxic signaling and eventual neuronal loss. We assessed these brain pathologies in aged canines and found an increase in the number of glial cells, both astrocytes and microglia, and the activation of astrocytes indicative of neuroinflammation. A rise in the aggregated protein Aβ_1-42_ and hyperphosphorylated tau, at Threonine 181 and 217, in the cortical brain regions of aging canines is seen. We then asked if any of these aged canines had CCD utilizing the only current diagnostic, owner questionnaires, verifying positive or severe CCD had pathologies of gliosis and accumulation of Aβ_1-42_ like their aged matched controls. However uniquely the CCD dogs had P-tau at T217. Therefore, this phosphorylation site of tau at threonine 217 may be a predictor for CCD.

## 1. Introduction

Age is the number one risk factor for cognitive decline in canines and humans; however, the etiology is not known. Therefore, it is thought that a clearer understanding of the neuropathogenesis of aging canine brains is essential for future the development of diagnostic tests and subsequently facilitate timely intervention. Studying aging dogs with and without canine cognitive dysfunction (CCD) syndrome shows promise for a better understanding of human age-related neurodegeneration as well. Canine cognitive dysfunction is considered an age-related disease, with the prevalence ranging from 14-35% of the senior canine population, exponentially increasing with age [1; 2; 3]. In a two-year longitudinal study of 51 dogs over the age of 8 years, 33% of dogs with normal cognitive status progressed to mild cognitive impairment and 22% of dogs with mild cognitive impairment progressed to CCD [4]. In another study of 180 geriatric dogs (aged 11-16 years), 28% of dogs 11-12 years old and 68% of dogs 15-16 years old exhibited signs of cognitive impairment [3]. With the aging canine population steadily increasing, it is essential to study these aging dogs, especially given the implications and translational value for the human counterpart.

The aging canine brain, similar to humans, correlates with a rise in cognitive decline and neuronal pathology [5; 6; 7; 8]. Unlike transgenic laboratory rodent models commonly used to study aging and cognitive decline, dogs age and develop CCD naturally, therefore making canines an ideal avenue for studying brain aging and disease. One common pathology in brain aging is an increase in glial inflammation usually affecting the frontal cortex and hippocampus [9]. Neuroinflammation in the CNS induces two phenotypes of reactive or activated astrocytes called A1 and A2 phenotypes [10], the A1 phenotype is pro-inflammatory and can induce neuronal death. The A1 reactive phenotype has been shown to be increased through the activation of microglia via the release of proinflammatory chemokines and cytokines [11]. Like astrocytes, microglia play an important role in inflammatory responses and aged canines exhibit greater levels of the microglial marker Iba-1 in the hippocampus of adult canines [12; 13; 14].

The aggregation and deposition Aβ as extracellular plaques and hyperphosphorylation of the tau protein are common brain pathologies seen in humans and some canines with cognitive decline. Amyloid-β_1-42_ was found to be present in the form of insoluble plaques in the cerebral cortex in humans and cognitive impairment in aged canines has been strongly associated with the accumulation of AB_1-42_ [8; 15; 16]. The amino acid sequence of β-amyloid is identical between humans and dogs, [17] allowing use of similar reagents across species. The hyperphosphorylation of the tau protein is also commonly found in humans with AD and is recently discovered in CCD dogs as well [18]. Therefore, we investigated the occurrence of P-tau and AB_1-42_ in our aged canine brains, using multiple antibodies for phosphorylation of tau and techniques to verify the role of this important misfolded protein in the aging canine.

## 2. Materials and Methods

### 2.1 Sample Collection and Preparation

Forty-seven canine brains were obtained from privately owned dogs by owner consent after canines were euthanized and necropsied. The brains were collected at the James L. Voss Veterinary Teaching Hospital at Colorado State University. Canine brains were fixed in 10% neutral buffer formalin. After fixation, brains were processed using a tissue processor (Leica TP 1020) and embedded in paraffin wax using a tissue embedder (Leica EG 1160). Tissue blocks containing the frontal cortex and the hippocampus region were cut using a microtome (ThermoFisher HM1030) at 5µm and mounted on a charged slide. The details for euthanasia, age, and weights of the canines are listed in **Table 1**.

**Table 1.**
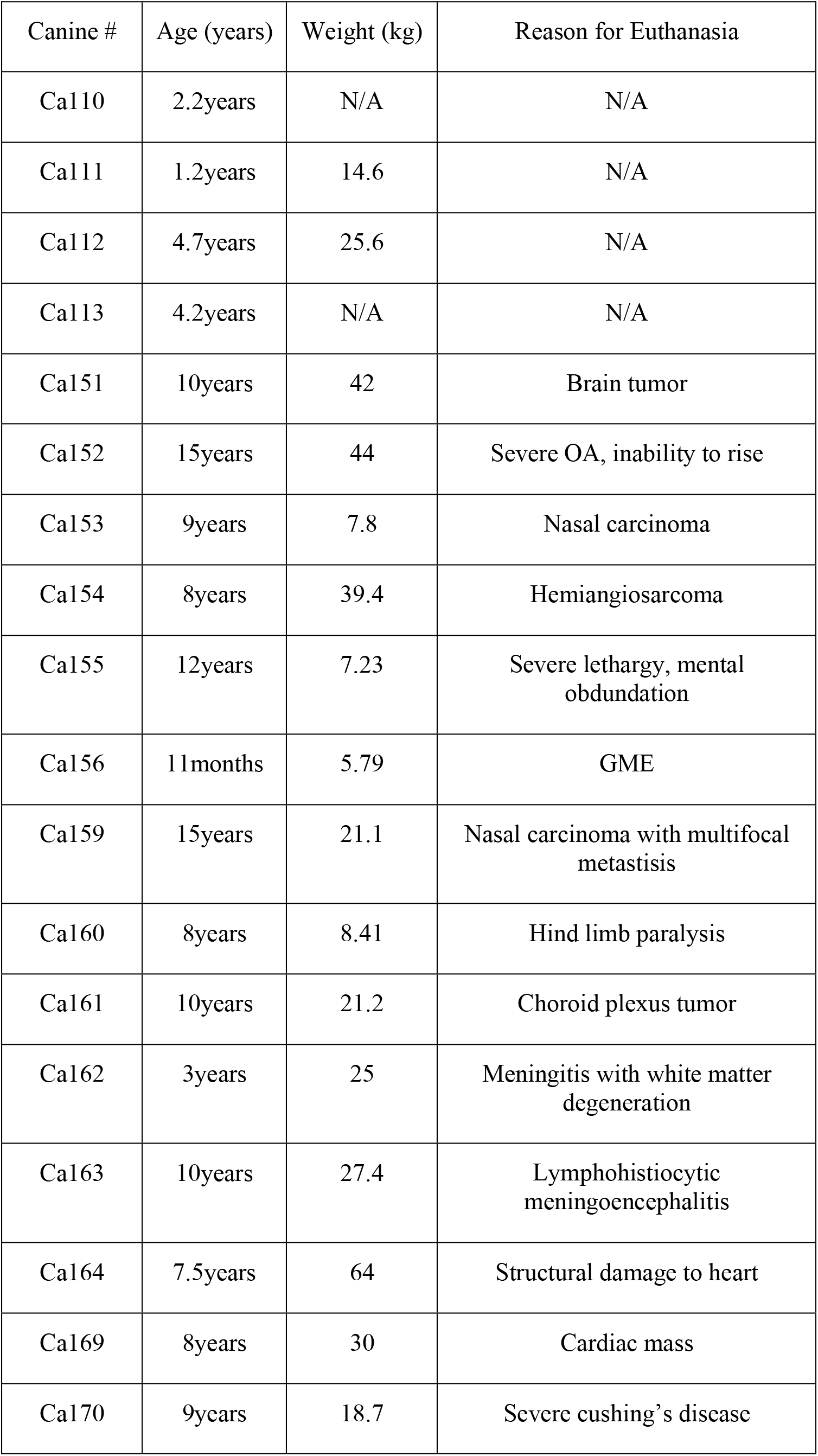
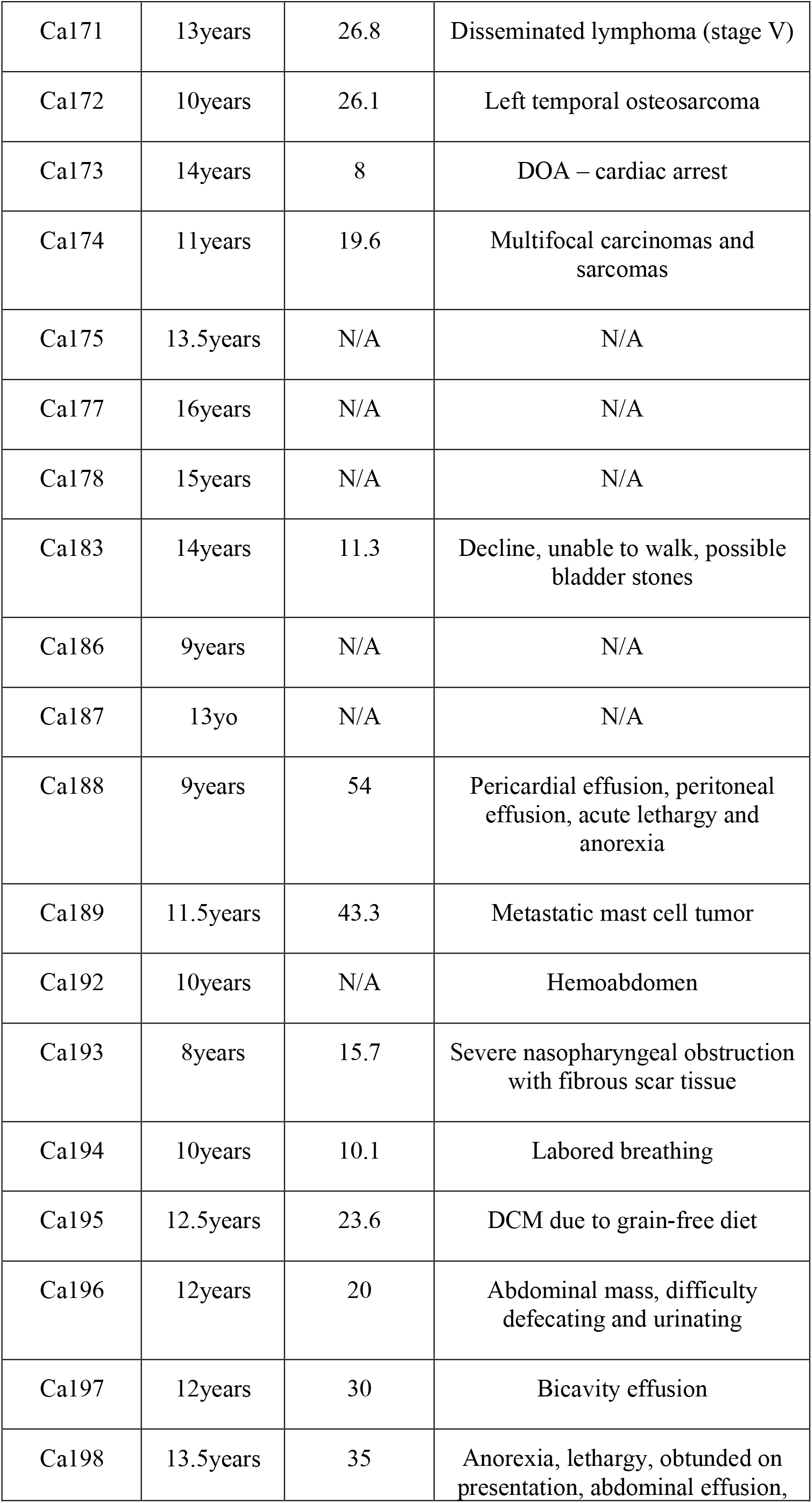
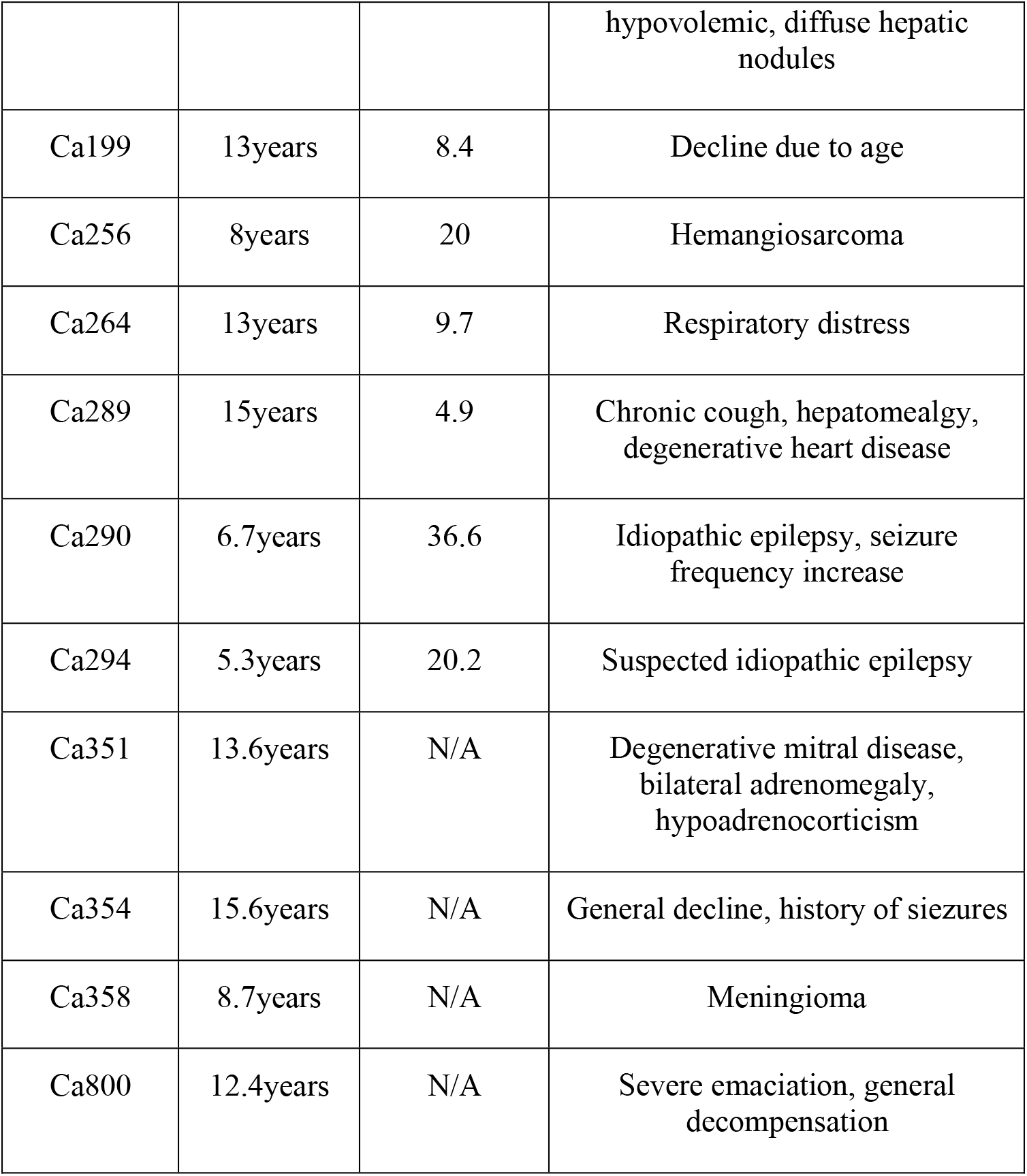
Aged canine cause of death, age and weight information.

| Canine # | Age (years) | Weight (kg) | Reason for Euthanasia |
| --- | --- | --- | --- |
| Ca110 | 2.2years | N/A | N/A |
| Ca111 | 1.2years | 14.6 | N/A |
| Ca112 | 4.7years | 25.6 | N/A |
| Ca113 | 4.2years | N/A | N/A |
| Ca151 | 10years | 42 | Brain tumor |
| Ca152 | 15years | 44 | Severe OA, inability to rise |
| Ca153 | 9years | 7.8 | Nasal carcinoma |
| Ca154 | 8years | 39.4 | Hemiangiosarcoma |
| Ca155 | 12years | 7.23 | Severe lethargy, mental obdundation |
| Ca156 | 11months | 5.79 | GME |
| Ca159 | 15years | 21.1 | Nasal carcinoma with multifocal metastasis |
| Ca160 | 8years | 8.41 | Hind limb paralysis |
| Ca161 | 10years | 21.2 | Choroid plexus tumor |
| Ca162 | 3years | 25 | Meningitis with white matter degeneration |
| Ca163 | 10years | 27.4 | Lymphohistiocytic meningoencephalitis |
| Ca164 | 7.5years | 64 | Structural damage to heart |
| Ca169 | 8years | 30 | Cardiac mass |
| Ca170 | 9years | 18.7 | Severe cushing's disease |
| Ca171 | 13years | 26.8 | Disseminated lymphoma (stage V) |
| Ca172 | 10years | 26.1 | Left temporal osteosarcoma |
| Ca173 | 14years | 8 | DOA – cardiac arrest |
| Ca174 | 11years | 19.6 | Multifocal carcinomas and sarcomas |
| Ca175 | 13.5years | N/A | N/A |
| Ca177 | 16years | N/A | N/A |
| Ca178 | 15years | N/A | N/A |
| Ca183 | 14years | 11.3 | Decline, unable to walk, possible bladder stones |
| Ca186 | 9years | N/A | N/A |
| Ca187 | 13yo | N/A | N/A |
| Ca188 | 9years | 54 | Pericardial effusion, peritoneal effusion, acute lethargy and anorexia |
| Ca189 | 11.5years | 43.3 | Metastatic mast cell tumor |
| Ca192 | 10years | N/A | Hemoabdomen |
| Ca193 | 8years | 15.7 | Severe nasopharyngeal obstruction with fibrous scar tissue |
| Ca194 | 10years | 10.1 | Labored breathing |
| Ca195 | 12.5years | 23.6 | DCM due to grain-free diet |
| Ca196 | 12years | 20 | Abdominal mass, difficulty defecating and urinating |
| Ca197 | 12years | 30 | Bicavity effusion |
| Ca198 | 13.5years | 35 | Anorexia, lethargy, obtunded on presentation, abdominal effusion, hypovolemic, diffuse hepatic nodules |
| Ca199 | 13years | 8.4 | Decline due to age |
| Ca256 | 8years | 20 | Hemangiosarcoma |
| Ca264 | 13years | 9.7 | Respiratory distress |
| Ca289 | 15years | 4.9 | Chronic cough, hepatomealgy, degenerative heart disease |
| Ca290 | 6.7years | 36.6 | Idiopathic epilepsy, seizure frequency increase |
| Ca294 | 5.3years | 20.2 | Suspected idiopathic epilepsy |
| Ca351 | 13.6years | N/A | Degenerative mitral disease, bilateral adrenomegaly, hypoadrenocorticism |
| Ca354 | 15.6years | N/A | General decline, history of siezures |
| Ca358 | 8.7years | N/A | Meningioma |
| Ca800 | 12.4years | N/A | Severe emaciation, general decompensation |

### 2.2 Immunohistochemical staining and Microscopy imaging analysis

Immunohistochemistry was used to visualize a variety of neurological antibodies including: GFAP (DAKO Cat. #20334), Iba1 (Abcam 5076), S100β (Abcam 41428), AT270 (Thermo Fisher MN1050), T217 (Abclonal AP1233) and Aβ_1-42_ (Thermo Fisher 44344). Sections were dewaxed in xylene and rehydrated through graded ethanol’s, before undergoing antigen retrieval in 0.01M sodium citrate for 20 mins at 95°C. Endoperoxides were removed by incubating sections in 0.30% hydrogen peroxide and water solution for 30 minutes. Blocking of non-specific labeling was performed with 10% hose serum in Tris-A in 2% BSA (2% bovine serum albumin (BSA) and 2% Triton-X in tris buffered saline (TBS)). Sections were incubated overnight at 4°C in primary antibodies diluted with Tris-A in 2% BSA. Sections were then washed with Tris-A in 2% BSA and incubated for an hour against their corresponding biotinylated secondary antibodies, goat anti-rabbit or horse anti-mouse (Vector labs). Slides were counterstained with hematoxylin, dehydrated through graded ethanol followed by xylene, mounted with mounting media and cover slipped with #1 cover glass.

Slides were visualized using Olympus BX51 microscope and Olympus DP70 camera and saved as virtual slide images at 20X objective. Quantitative analysis was performed using Olympus CellSens Dimension Desktop 3.1 using Count and Measure quantification application and mm^2^ region of interest was analyzed based on the brain area. Images were taken as virtual slide images at 40X objective to be analyzed on Olympus Cell Sense technology using Count and Measure quantification application. The frontal cortex and the hippocampus were chosen as the regions of interest.

### 2.3 Immunofluorescent Staining and Microscopy imaging analysis

Immunofluorescence was used to visualize co-expression of certain proteins in glial or neuronal cell types: C3(Abcam 181147) and S100β (Abcam 41428). Slides were visualized using BX63 fluorescence microscope equipped with a Hamamatsu ORCA-flash 4.0 LT CCD camera and collected using Olympus CellSens software version 3.1. Images were taken as virtual slide images at 10X objective to be analyzed on Olympus Cell Sense technology using Count and Measure quantification application.

## 3. Results

### 3.1 Increased glial inflammation in the frontal cortex and hippocampus of aged canines

Young (1-4 years of age) and aged (over 8 years of age) canine brains were stained and analyzed for numbers of glial markers including GFAP, S100β for astrocytes, and Iba1 for microglia in two key brain regions, frontal cortex and hippocampus. Representative images of the frontal cortex of a 3-, 9- and 12-year-old canine stained for GFAP (**Figure 1A-C**), Iba1 (**Figure 1E-G**) and S100β (**Figure 1I-K**) is shown. A significant increase is seen in S100β positive cells with a mean of 168.4 mm^2^ in aged canines compared to 92.67 mm^2^ in young dogs (**Figure 1L**) with the difference in means being 75.71±34.41. No significant difference was identified in the number of GFAP positive cells (**Figure 1D**) or Iba1 positive cells (**Figure 1H**), however a trend of an increase of Iba1 is detected in the frontal cortex.

**Figure 1.**
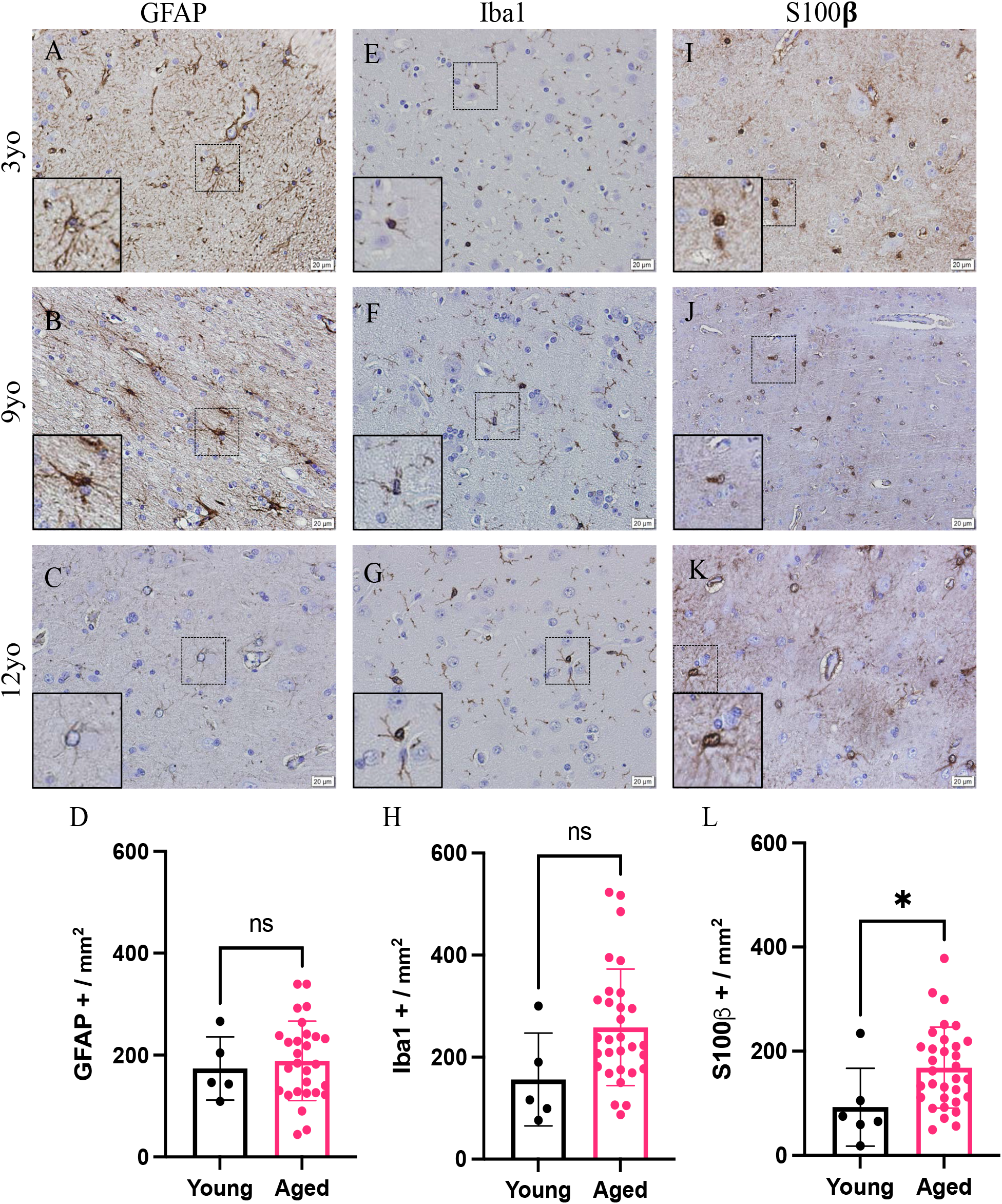
Increased glial reactivity in the frontal cortex of aging canines. Increase in astrocyte reactivity detected by increase in S100b+ (H, I, L; n=33) and microglia by IBA1+ cells (E, F, K; n=35) in the frontal cortex of aged canines when compared to young canines (D, G, J; n=38). No significant change was detected in GFAP+ cells. Scale bar = 20 µM Unpaired t-test, error bars = SEM, p< 0.05. p Values: D=0.6851, H=0.0673, L=0.0343

All available brains with a hippocampus, 13 of the 23 dogs analyzed in Figures 1 and 2, were stained with glial markers as well. Representative images of the hippocampus of 3-, 9- and 12-year-old canines stained for GFAP (Figure 2A-C), Iba1 (**Figure 2E-G**) and S100β (**Figure 2I-K**). Iba1 positive microglia significantly increased in the hippocampus of aged canines compared to young, with the mean of 274.4 mm^2^ in aged canines compared to a mean of 199.0mm^2^ for young canines. The difference between means is 75.4±27.04 (**Figure 2H**) p<0.05. A significant increase in S100β positive cells is also seen with a mean of 177.2 mm^2^ in aged canines compared to 69.5 mm^2^ in young canines (**Figure 2L**), the difference means being 107.7±35.4. No significant change, although a trend of an increase, was identified in the number of GFAP positive cells in the hippocampus (**Figure 2D**).

**Figure 2.**
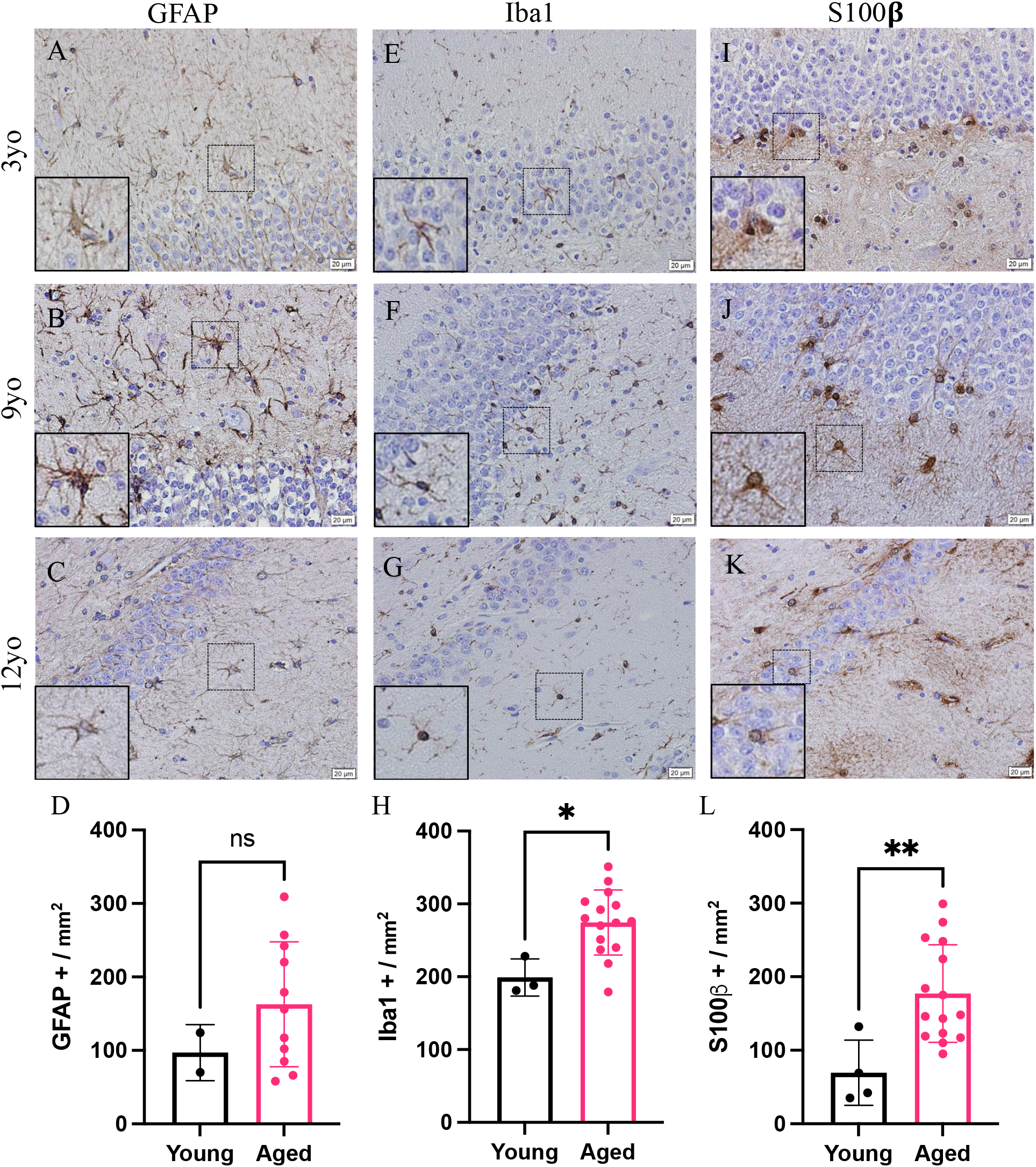
Increase glial reactivity in the hippocampus of aging canines. Increase in astrocyte reactivity detected by increase in S100β+ (J, K, L) and microglia by IBA1+ cells (F, G, H) in the hippocampus of aged canines when compared to young canines (A, E, I). No significant change was detected in GFAP+ cells. Scale bar = 20 µM Unpaired t-test, error bars = SEM, p< 0.05. p Values: D=0.2383, H=0.0010, L=0.0094

### 3.2 A1 astrocytes increase in the frontal cortex of aged canines

The frontal cortex of both young and aged canines was analyzed to identify co-localization of S100β and C3 staining using co-immunofluorescence. Representative images show colocalization of S100β and C3 staining of the aged canine in the frontal cortex (**Figure 3A, B**) and the hippocampus (**Figure 3C, D**). A significant increase in the percent of S100β to C3 was found (**Figure 3E**), with p < 0.05 and the difference between means being 22.94±10.95.

**Figure 3.**
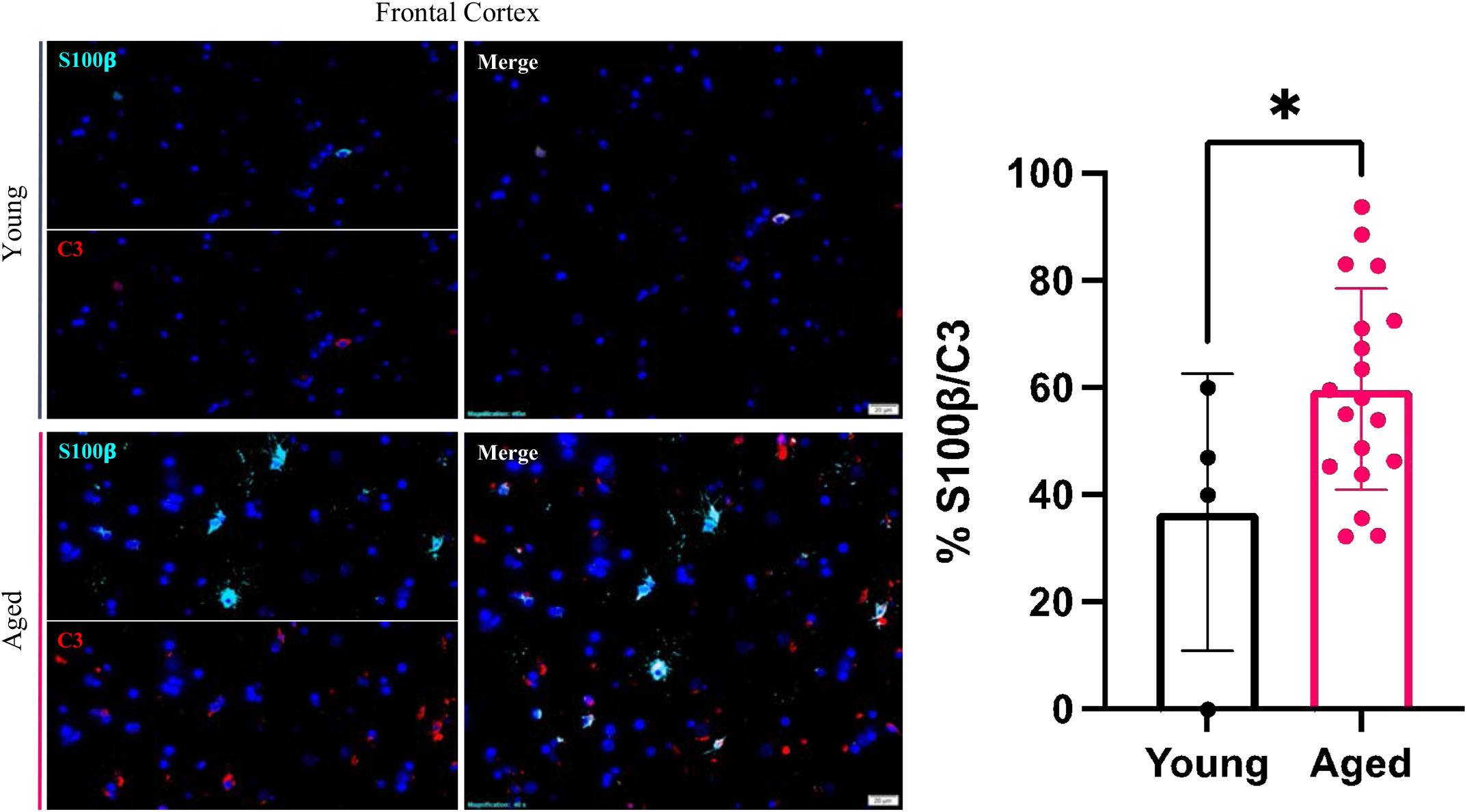
A1 reactive astrocytes increase within the frontal cortex and hippocampus of the aging canine. Pan astrocyte marker S100b (green) co-localization with C3 (red) reactivity reveals frontal cortex phenotypic switching in the aged canine (B) and the hippocampus (D), when compared to young canine brains, respectively (A, C). Quantification of A1 astrocytes in the frontal cortex (E, n=22). Scale bar = 20 µM Unpaired t-test, p=0.0486, error bars = SEM, p<0.05

### 3.3 Phosphorylated tau and amyloid beta plaques increase in the frontal cortex of aging canines

Representative images of hyperphosphorylation of tau and Aβ_1-42_ in the frontal cortex from a 3-, 9- and 12-year-old canines. Two different phosphorylation sites of tau (P-tau) were analyzed by immunohistochemistry including P-tau-Thr181 (Figure 4A-C) and P-tau-Thr217 (**Figure 4D-F**). An increase in extracellular and intracellular P-tau is found in the aging canine brains, 9- and 12-year old’s when compared to the young canine brain, 3-year-old. A possible formation of an extracellular neurofibrillary tangle with P-tau T217 antibody in the 9-year-old canine (see inset in **Figure 4E**) and intracellular P-tau aggregates in the 12-year-old canine (see inset **Figure 4F**). Amyloid-β_1-42_ aggregation was increased in the frontal cortex of 9- and 12-year-old canine compared to young dogs (**Figure G-I**).

**Figure 4.**
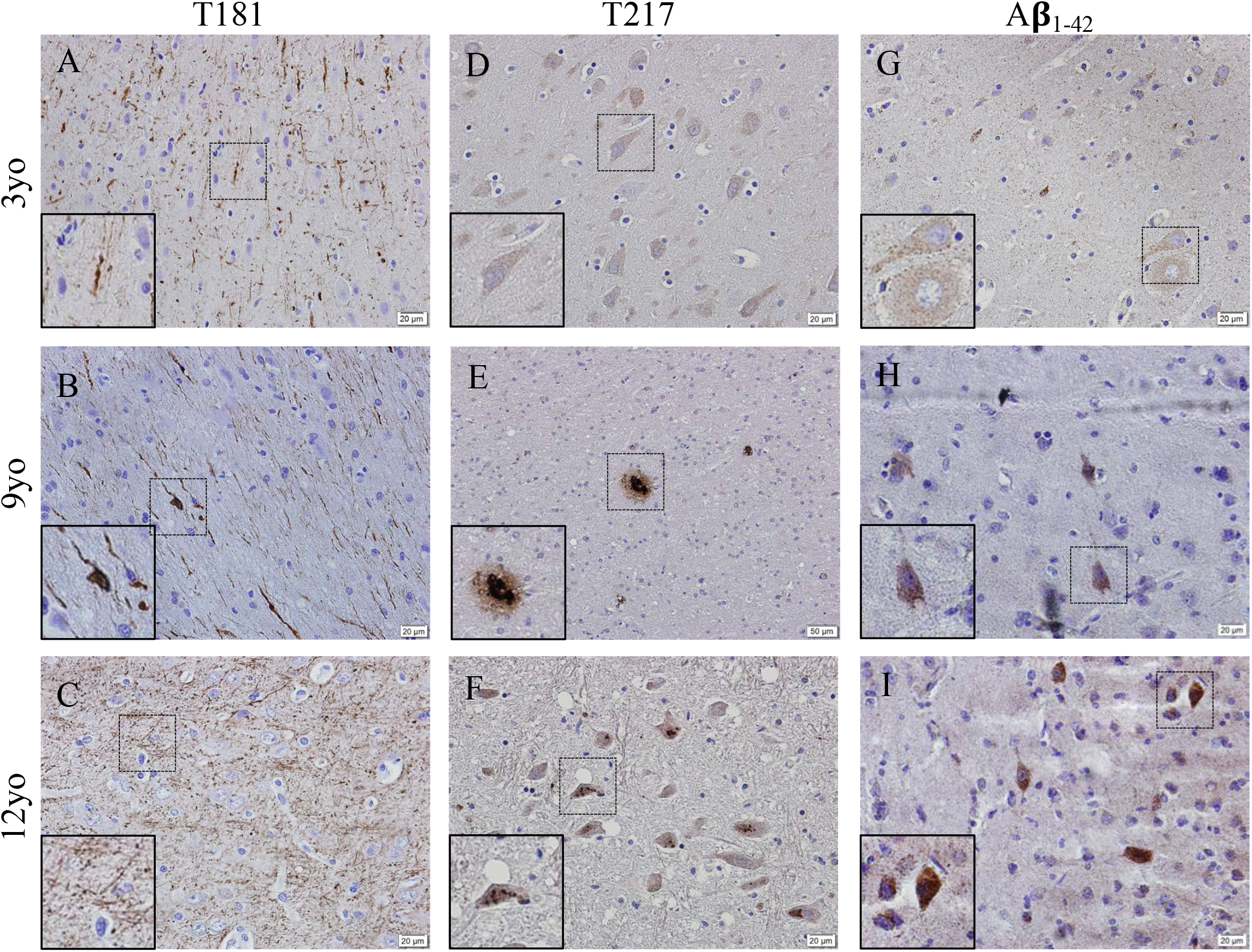
Hyperphosphorylation of Tau and Amyloid β_1-42_ accumulation in the frontal cortex of aging canines. Representative images of young (3 years of age; N= 5) and aged (8 years of age and older) canines are shown. Phosphorylated threonine 181 tau is increased at 9 (B) and 12 years of age (C). Phosphorylated threonine 217 tau is increased at 9 (E) and 12 years of age (F) with P-tau positive plaque like structure in (E, see inset) the 9-year-old dog and an intracellular phosphorylation in the 12-year-old canine (F, see inset). Aβ_1-42_ accumulates in the 9 (H) and 12 (I) year old dogs more than the young 3-year-old. Scale bar = 20µM

### 3.4 Questionnaires determined canine cognitive dysfunction in aging dogs and correlated to neuropathological results

The Canine Cognitive Dysfunction Rating Scale (CCDR) is based on a scale of 0-80 points, ranging from no abnormal behavior to severe behavioral disturbances [19]. The CCDR has a diagnostic accuracy of 99.3% in dogs. In accordance with previously published data, we used a score of 50 for the CCDR questionnaire as an optimal diagnostic threshold, meaning that we expected that dogs scoring more than 50 points were likely to have CCD [19]. The second questionnaire, Canine Dementia Scale (CADES), has been established as a validated screening tool for CCD, as well as a long-term assessment tool for progression and treatment efficacy [20]. Dogs were classified as having either mild (1-7), moderate (scores of 24-44) or severe cognitive impairment (scores of 45-95). Of the 19 owner questionnaires we received from the aging canines, 26% were positive by the CCDR scoring and 32% of the dogs were classified as having severe cognitive impairment from the CADES questionnaire (**Table 2**). Another 26% of the dogs were classified as either mild or moderate for cognitive impairment using the CADES (**Table 2**). Critically, aging canines with CCD scores of either positive by CCDR or severe with CADES showed an increase in gliosis or neuroinflammation and accumulation of both phosphorylated tau and Aβ_1-42_ (**Figure 6**). However, some of the aged canines with a negative score for CCDR or CADES were not distinguishable from the CCD brains (**Figure 6 E-H**).

**Table 2.**
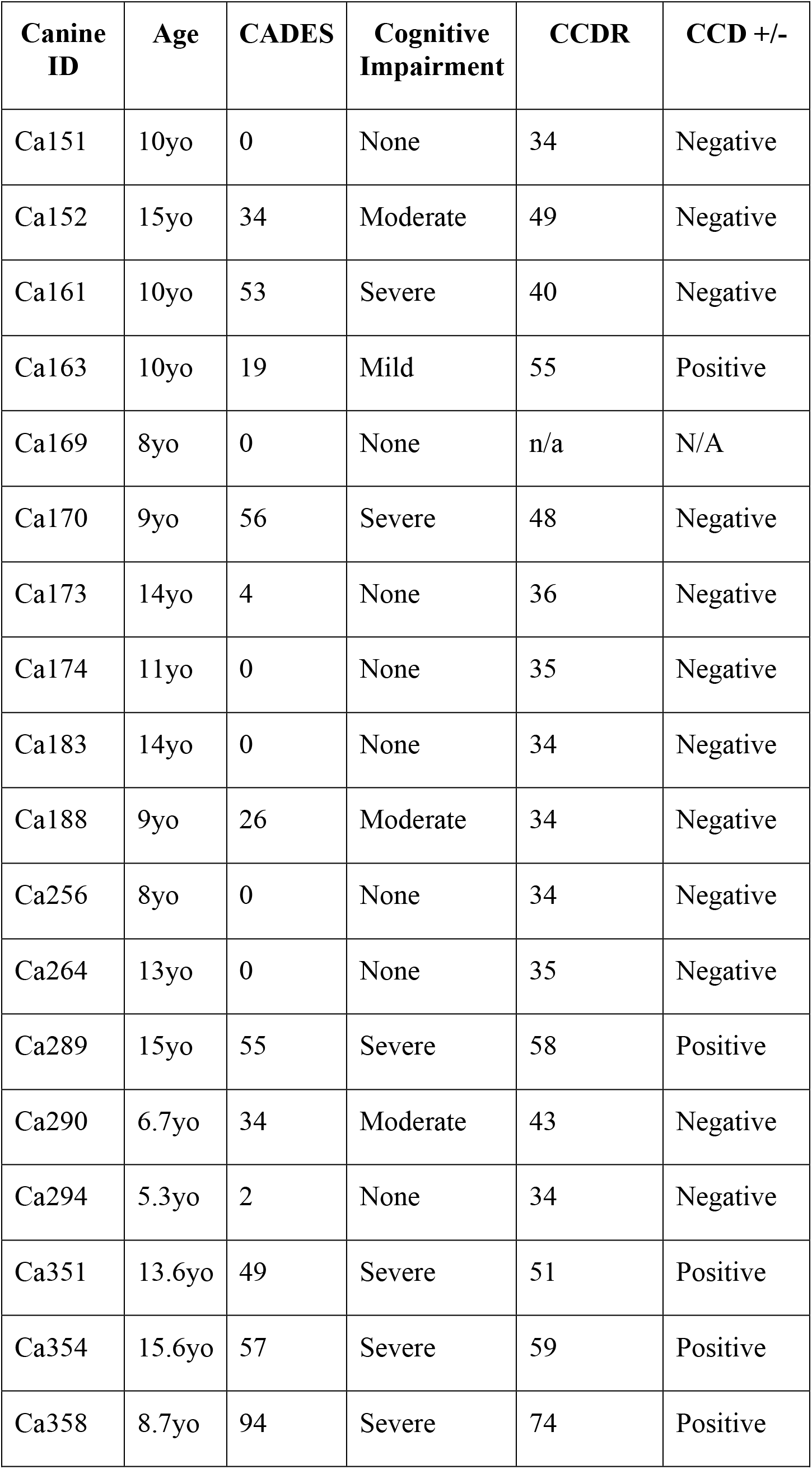
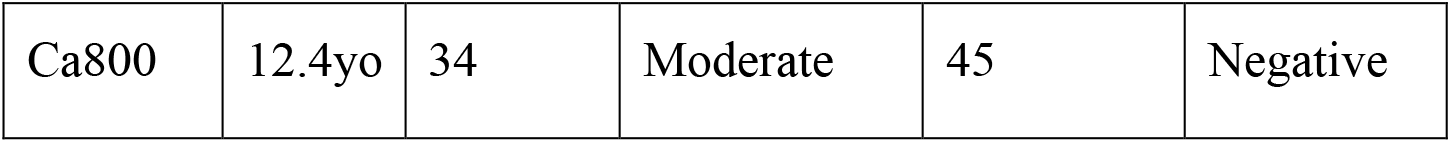
Scores of canines with CADES and CCDR owner questionnaires.

## 4. Discussion

Aged dogs with CCD show promise for modeling human age-related neurodegeneration, with classic hallmarks of Alzheimer’s disease (AD) and other AD related dementias (ADRDs) including CSF and brain Aβ, brain atrophy, and cognitive decline [4; 13; 14; 21; 22]. However, there is incomplete knowledge about how neuropathological progression is related to cognitive phenotype in canines. The aged canine is also able to share similar environmental factors as humans do, unlike lab research dogs or rodents, making them an ideal path to study the aging brain process. Here we quantify the amount of activated gliosis in aged canines as well as accumulation of the toxic proteins, Aβ_1-42_ and phosphorylation of tau (P-tau).

Microglia are essential for CNS homeostasis and play a role in age related neurodegenerative disease pathogenesis through the production of an inflammatory brain state [23; 24]. Our study is consistent with previous studies that aged canines exhibit greater levels of the Iba1 protein in the hippocampus than adult canines [13; 25] as well as in the canine cortical brain region (**Figures 1 and 2**). Further it is also known that with activation of microglial there is a progressive activation of astrocytes characterized by co-expression of C3 and S100β in AD human brains [11; 26]. Similar to microglia activated astrocytes produce numerous reactive and proinflammatory molecules in the AD brain [27]. The astrocyte marker S100β is also highly expressed by reactive astrocytes in close vicinity of beta amyloid deposits and immunotherapies for Aβ in dogs has shown to reduce S100β positive cells [28]. In this study we identify, C3 and S100β positive astrocytes significantly increased in aged canines indicating that as the canine ages activated astrocytes are accumulating (**Figure 3**).

Two hallmark features of AD in humans are the presence of neurofibrillary tangles, composed of abnormally hyperphosphorylated tau protein, and Aβ plaques. Canines share tight AD gene sequence homology with humans [29] and have alterations in amyloid precursor protein (APP) processing and tau immunoreactivity with age [30]. Aβ deposits also been observed in aging dog brains, with the load of Aβ often correlated with CCD clinical signs [31]. Similarly, phosphorylated tau protein has been detected in the aging dog brain, although with less frequency and density than human AD brains [14; 18]. Physiological tau is mostly confined to neuronal axons where it binds and helps stabilize microtubules. Microtubule binding and other tau functions are regulated by numerous serine, threonine and tyrosine phosphorylated sites within the protein. Cytoplasmic deposits of P-tau at Threonine181 have been identified in the frontal cortex of canines, but no neurofibrillary tangles (NFTs) have been identified [14]. In our study we revealed positivity of cells stained with P-tau Thr181 and Thr217 in cortices (**Figure 4B, C, E and F)** and hippocampus (**Figure 5B, C, E and F)** of aged canines compared to young canines, represented by Figure 4, 5A and D. Threonine 217 phosphorylation in the nine-year-old dog shows a plaque like structure or neurofibrillary tangle (NFT) (**Figure 4E**) while at 12-year-old dog has intracellular P-tau Thr217 staining (**Figure 4F**). These two differing cellular components of P-tau shows the variety that these of hyperphosphorylation these aging canines have similar to AD and ADRD patients [32; 33].

**Figure 5.**
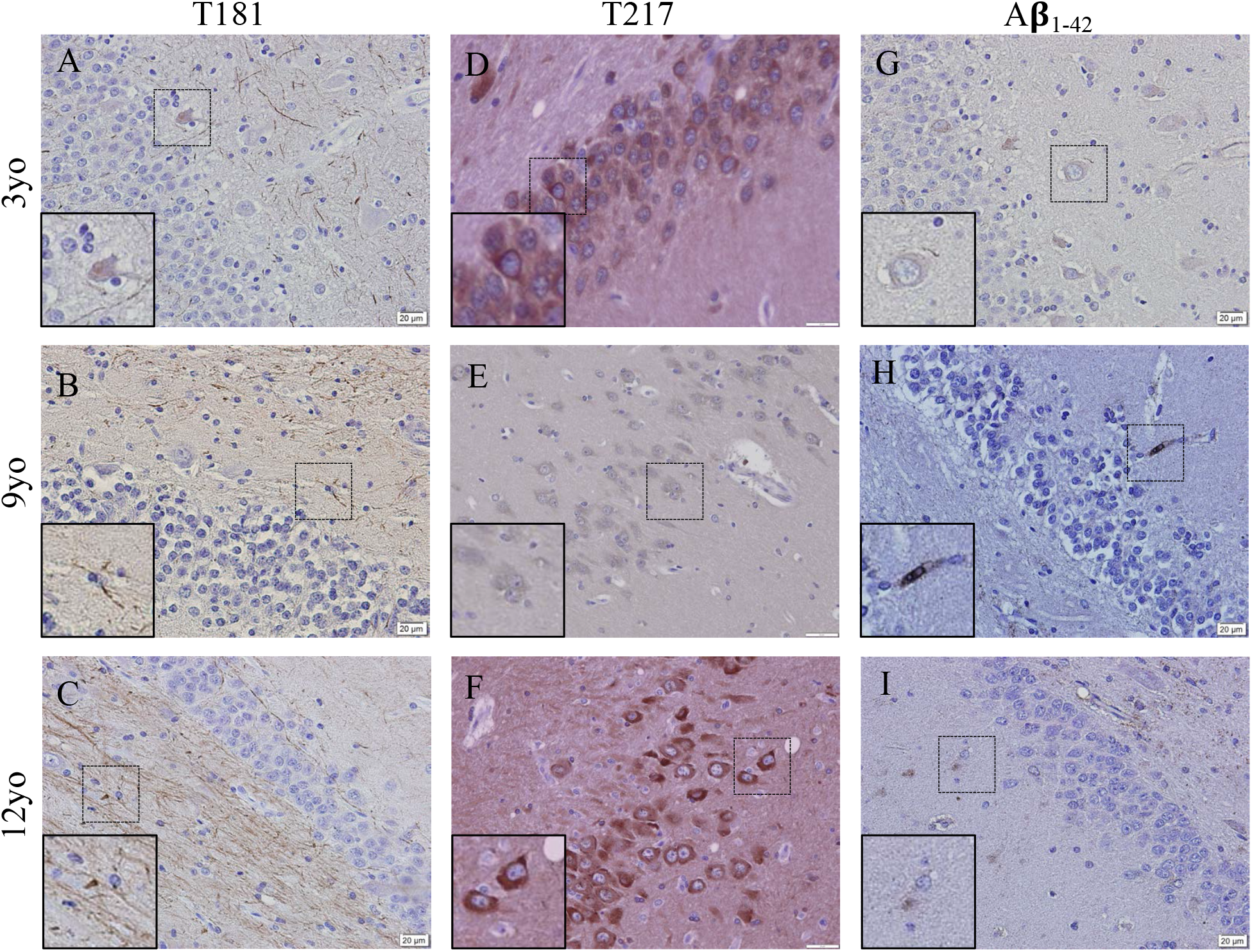
Gliosis and accumulation of misfolded proteins correlates to canines with positive cognitive dysfunction scores. (A) Aged canines with a CCDR/CADES score of being negative for CCD does show an increase in gliosis and Aβ_1-42_ accumulation but no P-tau at T-217. (B) Aged canines with a moderate or severe CCD score show gliosis and Aβ_1-42_ and P-tau positive cells. Scale bar = 20 µM

Lastly, it is critical that we are able to determine if these aging canines have CCD syndrome through the only current clinical diagnostic tool available, owner questionnaires [2; 34; 35; 36]. To determine this, we asked owners to complete both the CCDR and CADES questionnaires on necropsied dogs as none were diagnosed with CCD at the time of death (**Table 1**). All owners of the canines we have performed pathological analysis on were contacted and 19 or 40% of the surveys were returned for analysis (**Table 2**). However, pathology of the dogs with either severe criteria from the CADES questionnaire or CCDR positive for dementia show pathologies that were not always distinguishable from age matched canine brains. We found no significant change in the accumulation and aggregation of P-tau and Aβ (**Figure 6N &P**) in canines with CCD compared to aged matched canines without CCD in all of our canine brains. However, we did find, by using a morphological stain (H&E) a large number of punctate neurons (**Figure 6M**), indicative of apoptotic or dying neurons, in aged canines with CCD as compared to aged canines without CCD. The variability of our data indicated that there are a few limitations to our study that could explain our findings. The small sample size of owner questionnaires that we received may or may not be accurate as these were performed sometimes years after their dog had passed and we were unable to perform a medical review to rule out other causes of clinical disease, such as brain structural abnormalities or infection. Therefore, we are unsure if the lack of significance is due to the lack of power for the clinical questionaries, the lack of a full medical review or that there is no true difference between aging pathology with and without CCD clinical diagnosis. Future studies that are well powered with full medical review and timely owner questionnaires may allow us to attain more accurate results.

**Figure 6.**
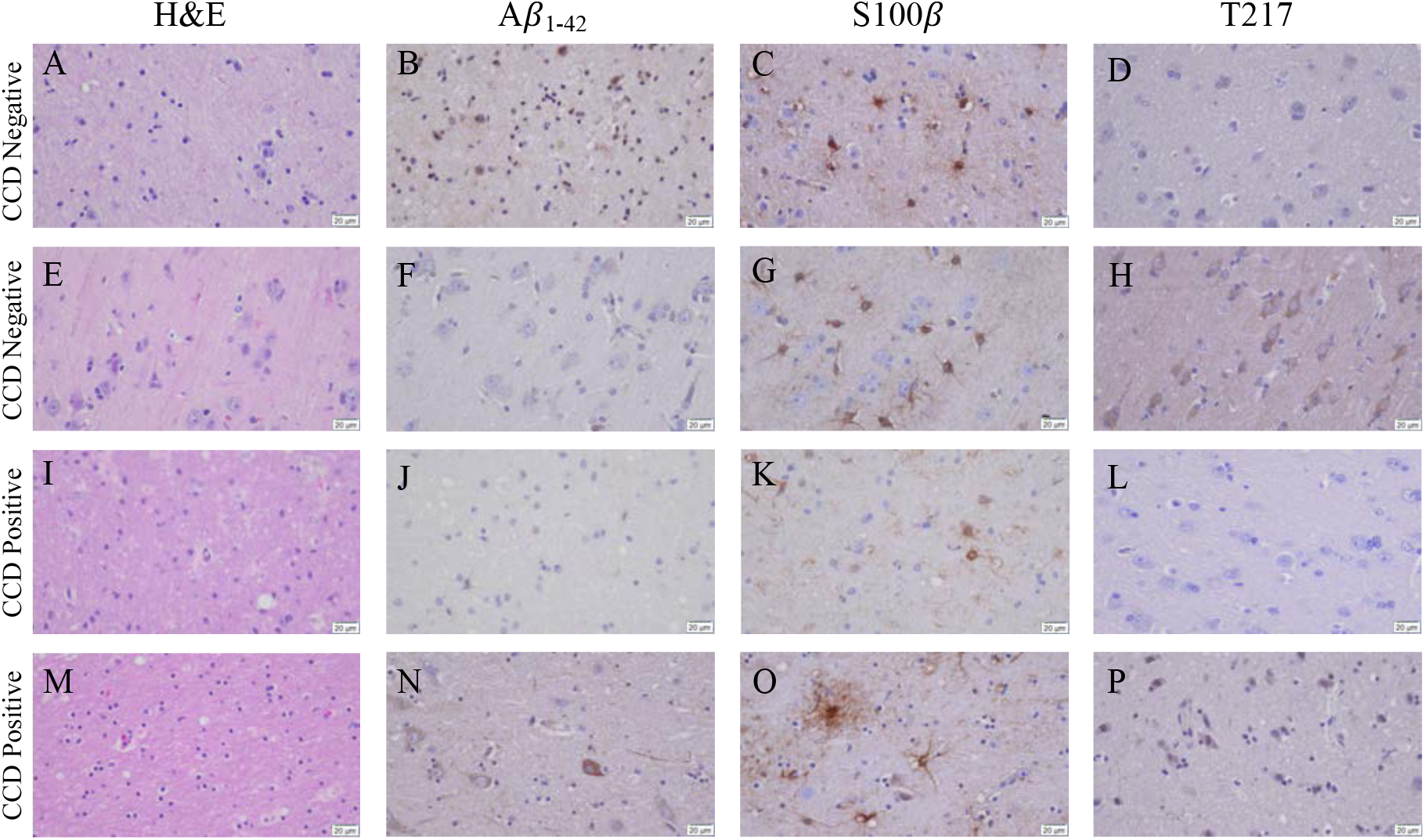
Gliosis and accumulation of misfolded proteins correlates to canines with positive cognitive dysfunction scores. (A) Aged canines with a CCDR/CADES score of being negative for CCD does show an increase in gliosis and Aβ_1-42_ accumulation but no P-tau at T-217. (B) Aged canines with a moderate or severe CCD score show gliosis and Aβ_1-42_ and P-tau positive cells. Scale bar = 20 µM

### 4.1 Conclusion

The aging brain and the factors that lead to susceptibility to diseases like cognitive decline is still unknown. To better understand this, we have studied thirty-eight naturally aged canine brains for pathological markers of glial inflammation, morphological changes, accumulation of amyloid β_1-42_ and hyperphosphorylation of tau. Both microglial and astrocyte number are increased, making this the first report to our knowledge detecting activation of reactive astrocytes defined by co staining with complement 3 (C3) and S100β in naturally aging canines, a similar phenotype found in Alzheimer’s disease laboratory rodent models and patients. We also detect Aβ_1-42_ accumulation and T181 and T217 for hyperphosphorylation of tau in most of the aged brain samples in this cohort. Following owner questionnaires, 19 of these aged dogs, 26% were positive for CCD based on the CCDR; however, there was no correlation between these biomarkers and CCD. However, future studies with timely clinical medical review and questionaries may allow us to determine differences between the two aging populations.

## 2 Author Contributions

J.A.M and S.M. conceived and designed the project. A.D.H, A.S.L, and M.L.R performed the pathology experiments. B.K. and L.M collected canine samples and owner questionnaires. A.D.H., S.M. and J.A.M contributed to data analysis. J.A.M wrote the manuscript and all authors reviewed and edited the manuscript.

## 4 Acknowledgments

We would like to acknowledge Dr. Savannah Rocha for their technical assistance for the co-immunofluorescence experiment.

## References

[1] G. Azkona, S. Garcia-Belenguer, G. Chacon, B. Rosado, M. Leon, and J. Palacio, Prevalence and risk factors of behavioural changes associated with age-related cognitive impairment in geriatric dogs. J Small Anim Pract 50 (2009) 87–91.

[2] H.E. Salvin, P.D. McGreevy, P.S. Sachdev, and M.J. Valenzuela, Under diagnosis of canine cognitive dysfunction: a cross-sectional survey of older companion dogs. Vet J 184 (2010) 277–81.

[3] J.C. Neilson, B.L. Hart, K.D. Cliff, and W.W. Ruehl, Prevalence of behavioral changes associated with age-related cognitive impairment in dogs. J Am Vet Med Assoc 218 (2001) 1787–91.

[4] T. Schutt, N. Toft, and M. Berendt, Cognitive Function, Progression of Age-related Behavioral Changes, Biomarkers, and Survival in Dogs More Than 8 Years Old. J Vet Intern Med 29 (2015) 1569–77.

[5] E. Head, H. Callahan, B.A. Muggenburg, C.W. Cotman, and N.W. Milgram, Visual-discrimination learning ability and beta-amyloid accumulation in the dog. Neurobiol Aging 19 (1998) 415–25.

[6] B.J. Cummings, and C.W. Cotman, Image analysis of beta-amyloid load in Alzheimer’s disease and relation to dementia severity. Lancet 346 (1995) 1524–8.

[7] E.C. Mormino, and K.V. Papp, Amyloid Accumulation and Cognitive Decline in Clinically Normal Older Individuals: Implications for Aging and Early Alzheimer’s Disease. J Alzheimers Dis 64 (2018) S633–S646.

[8] J.E. Rofina, A.M. van Ederen, M.J. Toussaint, M. Secreve, A. van der Spek, I. van der Meer, F.J. Van Eerdenburg, and E. Gruys, Cognitive disturbances in old dogs suffering from the canine counterpart of Alzheimer’s disease. Brain Res 1069 (2006) 216–26.

[9] T.G. Beach, R. Walker, and E.G. McGeer, Patterns of gliosis in Alzheimer’s disease and aging cerebrum. Glia 2 (1989) 420–36.

[10] D.P.Q. Clark, V.M. Perreau, S.R. Shultz, R.D. Brady, E. Lei, S. Dixit, J.M. Taylor, P.M. Beart, and W.C. Boon, Inflammation in Traumatic Brain Injury: Roles for Toxic A1 Astrocytes and Microglial-Astrocytic Crosstalk. Neurochem Res 44 (2019) 1410–1424.

[11] S.A. Liddelow, K.A. Guttenplan, L.E. Clarke, F.C. Bennett, C.J. Bohlen, L. Schirmer, M.L. Bennett, A.E. Munch, W.S. Chung, T.C. Peterson, D.K. Wilton, A. Frouin, B.A. Napier, N. Panicker, M. Kumar, M.S. Buckwalter, D.H. Rowitch, V.L. Dawson, T.M. Dawson, B. Stevens, and B.A. Barres, Neurotoxic reactive astrocytes are induced by activated microglia. Nature 541 (2017) 481–487.

[12] I.K. Hwang, C.H. Lee, H. Li, K.Y. Yoo, J.H. Choi, D.W. Kim, D.W. Kim, H.W. Suh, and M.H. Won, Comparison of ionized calcium-binding adapter molecule 1 immunoreactivity of the hippocampal dentate gyrus and CA1 region in adult and aged dogs. Neurochem Res 33 (2008) 1309–15.

[13] M. Ozawa, J.K. Chambers, K. Uchida, and H. Nakayama, The Relation between canine cognitive dysfunction and age-related brain lesions. J Vet Med Sci 78 (2016) 997–1006.

[14] T. Smolek, A. Madari, J. Farbakova, O. Kandrac, S. Jadhav, M. Cente, V. Brezovakova, M. Novak, and N. Zilka, Tau hyperphosphorylation in synaptosomes and neuroinflammation are associated with canine cognitive impairment. J Comp Neurol 524 (2016) 874–95.

[15] J. Barnes, P. Cotton, S. Robinson, and M. Jacobsen, Spontaneous Pathology and Routine Clinical Pathology Parameters in Aging Beagle Dogs: A Comparison With Adolescent and Young Adults. Vet Pathol 53 (2016) 447–55.

[16] M. Pugliese, J.L. Carrasco, C. Andrade, E. Mas, J. Mascort, and N. Mahy, Severe cognitive impairment correlates with higher cerebrospinal fluid levels of lactate and pyruvate in a canine model of senile dementia. Prog Neuropsychopharmacol Biol Psychiatry 29 (2005) 603–10.

[17] E.M. Johnstone, M.O. Chaney, F.H. Norris, R. Pascual, and S.P. Little, Conservation of the sequence of the Alzheimer’s disease amyloid peptide in dog, polar bear and five other mammals by cross-species polymerase chain reaction analysis. Brain Res Mol Brain Res 10 (1991) 299–305.

[18] A. Abey, D. Davies, C. Goldsbury, M. Buckland, M. Valenzuela, and T. Duncan, Distribution of tau hyperphosphorylation in canine dementia resembles early Alzheimer’s disease and other tauopathies. Brain Pathol 31 (2021) 144–162.

[19] H.E. Salvin, P.D. McGreevy, P.S. Sachdev, and M.J. Valenzuela, The canine cognitive dysfunction rating scale (CCDR): a data-driven and ecologically relevant assessment tool. Vet J 188 (2011) 331–6.

[20] A. Madari, J. Farbakova, S. Katina, T. Smolek, P. Novak, T. Weissova, M. Novak, and N. Zilka, Assessment of severity and progression of canine cognitive dysfunction syndrome using the CAnine DEmentia Scale (CADES). Applied Animal Behaviour Science 171 (2015) 138–145.

[21] C.H. Yu, G.S. Song, J.Y. Yhee, J.H. Kim, K.S. Im, W.G. Nho, J.H. Lee, and J.H. Sur, Histopathological and immunohistochemical comparison of the brain of human patients with Alzheimer’s disease and the brain of aged dogs with cognitive dysfunction. J Comp Pathol 145 (2011) 45–58.

[22] M. Pugliese, M.C. Geloso, J.L. Carrasco, J. Mascort, F. Michetti, and N. Mahy, Canine cognitive deficit correlates with diffuse plaque maturation and S100beta (-) astrocytosis but not with insulin cerebrospinal fluid level. Acta Neuropathol 111 (2006) 519–28.

[23] A. Sobue, O. Komine, Y. Hara, F. Endo, H. Mizoguchi, S. Watanabe, S. Murayama, T. Saito, T.C. Saido, N. Sahara, M. Higuchi, T. Ogi, and K. Yamanaka, Microglial gene signature reveals loss of homeostatic microglia associated with neurodegeneration of Alzheimer’s disease. Acta Neuropathol Commun 9 (2021) 1.

[24] W.Y. Wang, M.S. Tan, J.T. Yu, and L. Tan, Role of pro-inflammatory cytokines released from microglia in Alzheimer’s disease. Ann Transl Med 3 (2015) 136.

[25] B.B. Thomsen, C. Madsen, K.T. Krohn, C. Thygesen, T. Schutt, A. Metaxas, S. Darvesh, J.S. Agerholm, M. Wirenfeldt, M. Berendt, and B. Finsen, Mild Microglial Responses in the Cortex and Perivascular Macrophage Infiltration in Subcortical White Matter in Dogs with Age-Related Dementia Modelling Prodromal Alzheimer’s Disease. J Alzheimers Dis 82 (2021) 575–592.

[26] T. Wu, B. Dejanovic, V.D. Gandham, A. Gogineni, R. Edmonds, S. Schauer, K. Srinivasan, M.A. Huntley, Y. Wang, T.M. Wang, M. Hedehus, K.H. Barck, M. Stark, H. Ngu, O. Foreman, W.J. Meilandt, J. Elstrott, M.C. Chang, D.V. Hansen, R.A.D. Carano, M. Sheng, and J.E. Hanson, Complement C3 Is Activated in Human AD Brain and Is Required for Neurodegeneration in Mouse Models of Amyloidosis and Tauopathy. Cell Rep 28 (2019) 2111–2123 e6.

[27] F.L. Heppner, R.M. Ransohoff, and B. Becher, Immune attack: the role of inflammation in Alzheimer disease. Nat Rev Neurosci 16 (2015) 358–72.

[28] M. Neus Bosch, M. Pugliese, C. Andrade, J. Gimeno-Bayon, N. Mahy, and M.J. Rodriguez, Amyloid-beta immunotherapy reduces amyloid plaques and astroglial reaction in aged domestic dogs. Neurodegener Dis 15 (2015) 24–37.

[29] M.J. Sharman, S.H. Moussavi Nik, M.M. Chen, D. Ong, L. Wijaya, S.M. Laws, K. Taddei, M. Newman, M. Lardelli, R.N. Martins, and G. Verdile, The Guinea Pig as a Model for Sporadic Alzheimer’s Disease (AD): The Impact of Cholesterol Intake on Expression of AD-Related Genes. PLoS One 8 (2013) e66235.

[30] K. Bates, R. Vink, R. Martins, and A. Harvey, Aging, cortical injury and Alzheimer’s disease-like pathology in the guinea pig brain. Neurobiol Aging 35 (2014) 1345–51.

[31] E. Head, V. Pop, F. Sarsoza, R. Kayed, T.L. Beckett, C.M. Studzinski, J.L. Tomic, C.G. Glabe, and M.P. Murphy, Amyloid-beta peptide and oligomers in the brain and cerebrospinal fluid of aged canines. J Alzheimers Dis 20 (2010) 637–46.

[32] C.M. Moloney, V.J. Lowe, and M.E. Murray, Visualization of neurofibrillary tangle maturity in Alzheimer’s disease: A clinicopathologic perspective for biomarker research. Alzheimers Dement 17 (2021) 1554–1574.

[33] M. Wennstrom, S. Janelidze, K.P.R. Nilsson, B. Netherlands Brain, G.E. Serrano, T.G. Beach, J.L. Dage, and O. Hansson, Cellular localization of p-tau217 in brain and its association with p-tau217 plasma levels. Acta Neuropathol Commun 10 (2022) 3.

[34] M. Ozawa, M. Inoue, K. Uchida, J.K. Chambers, Y. Takeuch, and H. Nakayama, Physical signs of canine cognitive dysfunction. J Vet Med Sci 81 (2019) 1829–1834.

[35] H.E. Salvin, P.D. McGreevy, P.S. Sachdev, and M.J. Valenzuela, The canine sand maze: an appetitive spatial memory paradigm sensitive to age-related change in dogs. J Exp Anal Behav 95 (2011) 109–18.

[36] M.L. O’Brian, M.E. Herron, A.M. Smith, and T.K. Aarnes, Effects of a four-week group class created for dogs at least eight years of age on the development and progression of signs of cognitive dysfunction syndrome. J Am Vet Med Assoc 259 (2021) 637–643.

